# A model for maxilloturbinate morphogenesis in seals

**DOI:** 10.1101/2023.12.15.571824

**Authors:** Jonathan E. Kings, Lars P. Folkow, Øyvind Hammer, Signe Kjelstrup, Matthew J. Mason, Fengzhu Xiong, Eirik G. Flekkøy

## Abstract

The nasal cavities of mammals contain the maxilloturbinate bones, which are involved in respiratory heat and humidity regulation. The maxilloturbinates of Arctic seals develop into particularly elaborate labyrinthine patterns, which are well adapted to retain heat and moisture from exhaled gas. These structures develop prenatally and continue to grow postnatally. The developmental mechanism of labyrinthine patterning is unknown. Here we report a model of maxilloturbinate pattern formation in prenatal and juvenile seals based on a simple algorithmic description and three key parameters: target turbinate porosity, characteristic ossification time scale, and typical gestation time scale. Under a small set of geometrical and physical rules, our model reproduces key features of the patterns observed in grey and harp seal turbinates, and even in the less complex monk seal turbinates. To validate our model, we measure complexity, hydraulic diameter, backbone fractal dimension, and Horton-Strahler statistics for a rigorous quantitative comparison with actual tomograms of grey and harp seal skull specimens. Our model closely replicates the structural development of seal turbinates in these respects. Labyrinthine maxilloturbinate development likely requires the ability for neighbouring bone branches to detect and avoid each other through the mechanosensing of shear stresses from amniotic fluid and air flow.

## 1 Introduction

Maxilloturbinates in polar and subpolar phocid seal species (Fig. 1) develop into highly intricate dendritic patterns (Folkow et al., 1988; Mason et al., 2020; Negus, 1958; Van Valkenburgh et al., 2011), which play an important role in heat and water homeostasis (Flekkøy et al., 2023; Folkow & Blix, 1987; Huntley et al., 1984). The bone structure is connected dorsolaterally in the rostral part of the nasal cavity, rooted on the maxillary bone on either side, with branches more densely packed towards the middle and tapering off rostrally and caudally (Folkow et al., 1988; Mason et al., 2020). The single, ridge-like origin of the maxilloturbinate in the rostral part of the nasal cavity typically shortens and divides into several smaller roots in the posterior region.

**Figure 1:**
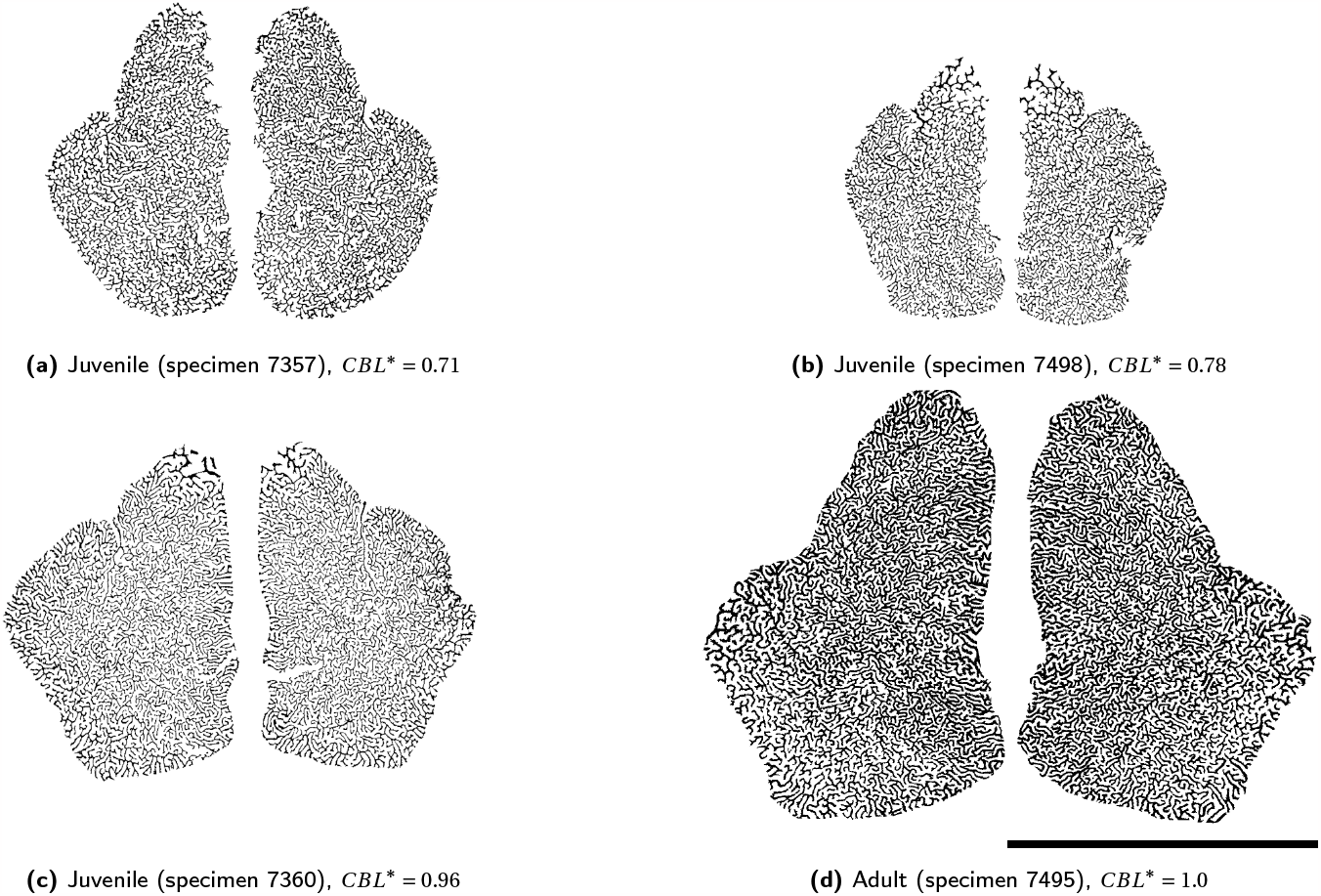
Harp seal maxilloturbinate tomograms ordered from youngest to oldest. The tomograms are reoriented through the densest part of the maxilloturbinate masses, and the specimens’ relative ages are based on measured condylobasal lengths, CBL. Maxilloturbinate bones shown in black, airways in white. Enclosing nasal cavity walls not shown. *CBL*^∗^ values shown are relative to adult size (*CBL*^∗^ *=* 1). Scale bar 50 mm.

Although pinniped turbinate development *per se* has not yet been studied extensively (Mason et al., 2020), we are assuming here that mechanisms uncovered from developmental studies across other mammal groups are conserved in pinniped development, as summarised below.

In pigs (*Sus scrofa*), longitudinal maxilloturbinate growth involves endochondral ossification which extends in the rostral direction (Martineau-Doizé et al., 1992). Transversally, the maxilloturbinate branches are shown to grow appositionally in both fruit bats (*Rousettus leschenaultii*) (Smith et al., 2021) and pigs (Martineau-Doizé et al., 1992). Growth at the tips of the branches is consistent with early branching followed by postnatal branch length growth, proposed for grey seals by Mason et al. (2020) on the basis of a comparison of neonatal and adult tomograms.

Growth of branched turbinates is bilaterally asymmetrical beyond the initial main branches (Hillenius, 1992; Negus, 1958). Their patterns are apparently not identical between any two individuals of the same species (Van Valkenburgh et al., 2011), which strongly suggests that the growth pattern is not rigidly pre-programmed. This aligns with the conclusions of Johnston et al. (2022) that biological patterns tend to be at least partly algorithmically encoded rather than entirely pre-programmed.

These key features of the pattern can shed light on its underlying developmental mechanisms. Morphological patterns in developing organisms often emerge as a consequence of physical and chemical interactions between the environment and cells, following a set of simple and limited local rules, making them productive subjects to study with mathematical models (Bard, 1981; Maini, 2004). In non-biological systems, model-based approaches have helped uncover an understanding of the underlying mechanisms of pattern formation, such as frictional fingering labyrinthine patterns in granular fluid dynamics (Olsen et al., 2019; Sandnes et al., 2007) and ferrofluid labyrinths in ferrohydrodynamics (Langer et al., 1992). A prominent example in biology is the class of “Turing” models recapitulating the formation of periodic biological patterns via reaction-diffusion (Turing, 1952) and related instabilities (Hiscock & Megason, 2015), such as in fingerprints (Glover et al., 2023) and zebra stripes (Murray, 1988). We chose here an approach that involves a direct and continued role of mechanics in the development of turbinate structures, akin to pattern formation models that highlight the role of mechanical forces in directly shaping the tissue, such as in epithelial invagination during branching morphogenesis in lungs (Miura, 2015). Using this approach, we introduce below a novel geometrical growth model to simulate seal maxilloturbinate pattern formation, with the goal of reproducing the most important structural features as simply as possible. Having identified the key requirements of the model, we are in a position to suggest biological mechanisms which are likely to underlie seal maxilloturbinate development.

## 2 Methods

While the maxilloturbinate labyrinthine patterns vary gradually along the direction of the flow, the cross-sectional patterns in the transverse plane remain consistent and encapsulate all key structural features. Therefore, to effectively model maxilloturbinate development, we focused on a two-dimensional cross-section in the transverse plane, based on the following six assumptions:

1. It grows within a space representing the nasal cavity, which becomes confining.
2. Growth occurs only at the tips of the structure, and branching may occur only via forking. We do not include any bone resorption in our model.
3. The cross-sectional structure has an elastic response that resists bending and locally minimizes its curvature.
4. As the structure grows, older parts become more rigid than new ones. The increase in rigidity occurs over a time scale comparable to the duration of the prenatal growth.
5. A self-avoidance mechanism keeps the branches from growing into each other.
6. All branches are established prenatally, with postnatal growth being limited solely to the elongation of preexisting branches.

Our model describes an implementation of these constraints by means of overdamped relaxation dynamics (Ling et al., 2016). The forces that drive this dynamical process are designed to reproduce the geometric effects of the underlying biological mechanisms but not necessarily to simulate the mechanisms involved. This is particularly true for the force that creates self-avoidance in the model.

In a two-dimensional transversal plane, the turbinate can be seen as a bony tree whose branches display slight thickness variations. Abstracting away the thickness of the bone branches, we define the geometrical backbone of the maxilloturbinate as the set of thin lines in the centre of these branches, i.e. an infinitely thinned version of the bone’s core. We thereby model the turbinate’s backbone as a one-dimensional elastic tree of connected nodes embedded within a two-dimensional plane, with no inherent branch thickness.

The growing branches are constrained within a circular boundary of maximum radius *B* corresponding to the nasal cavity (Fig. 2). The node backbone tree is constructed iteratively by alternating two separate steps: a *relaxation* step and a *growth* step.

**Figure 2:**
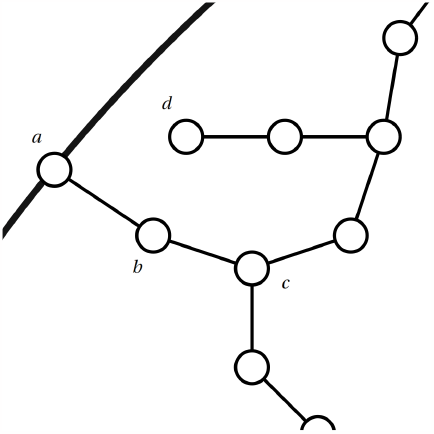
We model the two-dimensional backbone of a maxilloturbinate as a series of connected nodes. The nodes are constrained within a closed circular boundary representing the nasal cavity wall (thick line) of radius *B* . The first node (*a*) is fixed to the boundary. Neighbour nodes (*a* and *b, b* and *c*, etc.) are kept within a certain distance *r*_0_ of each other by means of an overdamped spring force along the connection. Non-neighbour nodes (*b* and *d, a* and *c*, etc.) repel each other with a characteristic minimum distance 2*r*_0_. New nodes grow from branch tips (e.g. at *d*). Except for *a*, all nodes are repelled from the boundary.

### Relaxation step

The relaxation step yields the position **r**_*i*_ of node *i* ≠ 0 at time *t* as

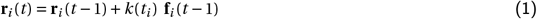

where *k*(*t*_*i*_) is a stifness coefficient decaying to 0 in characteristic rigidifying time scale *τ* with time *t*_*i*_ that has passed since the creation of the node *i*, as 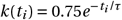.

The force **f**_*i*_ acting on each node is given by the sum of a repulsive force 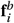 pushing nodes away from the boundary and node-node interaction forces 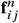 and 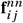 acting respectively from neighbour and non-neighbour nodes, *j* :

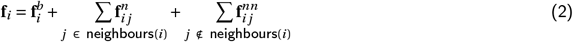

These interaction potentials act to keep neighbour nodes such as *b* and *c* in Fig. 2 within a characteristic distance *r*_0_ of each other and to keep non-neighbour nodes such as *b* and *d* in Fig. 2 away from each other:

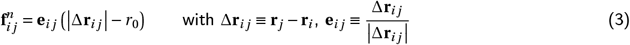

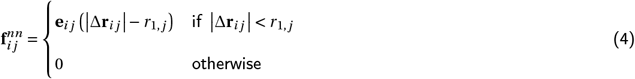

The non-neighbour interaction potential is a purely repulsive force and keeps non-neighbour nodes a distance of at least *r*_1, *j*_ from each other, defined as *r*_1, *j*_ *=* 2*r*_0_ or ∆**r**_*jh*_ (where *h* is the furthest neighbour of node *j*), whichever is greater, to ensure self-avoidance between different pieces of the node tree, thus effectively closing gaps in the chain. The chosen value of *r*_0_ only sets the numerical scale over which the pattern evolves, and as such does not influence the structure itself. Node *i =* 0 remains attached to the boundary throughout.

The boundary radius *b* and the labyrinth of cross-sectional bone area *A*_*b*_ define the porosity, i.e. the proportion of air within the outer circumference, *φ =* 1 − *A*_*b*_ /(*πb*^2^). Soft tissue covering the maxilloturbinates can be considered to be included within the area *A*_*b*_ . Since the model only describes the backbone of the turbinate section and a constant branch thickness is assumed, *A*_*b*_ is proportional to the total backbone length. In order to attain the target porosity *φ*_target_ *=* 0.75 observed across Arctic seal tomograms, we replace *b* by *b*^*′*^ where *φ*_target_ *=* 1 − *A*_*b*_ /(*πb*^*′*2^). Using the above equation for *φ* to replace *A*_*b*_ by *A*_*b*_ *=* (1 −*φ*)*πb*^2^, we may solve for *b*^*′*^ to get

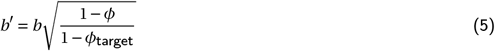

which is the new boundary radius, and increases up to a maximum of *B* . In our model the confining cavity therefore grows as a result of the developing turbinate system. Biologically, the causal relationship must be inverted, with turbinates developing within the enclosing cavity. However, this is simply an implementation detail: from a mathematical standpoint, identical results would be obtained if the sequence were reversed, with the enclosing boundary updated first and the turbinate as a result.

### Growth step

In order to model appositional growth, insertion of a new node is done either by extending a branch tip (e.g. from node *d* in Fig. 2) thus lengthening the branch, or by creating a fork at the tip of an existing branch. These node insertions throw the system off-equilibrium locally, requiring this growth step to be followed by 5 relaxation steps in order to restore a semi-equilibrium state before adding further nodes. The parent node, where growth takes place, is selected with a probability inversely proportional to local density. This gives priority to growth in low-density regions of the node tree, preventing high-density regions from becoming even denser. The position of the new node is chosen randomly on a circle of radius *r*_2_ around the parent node. The distance *r*_2_ is chosen as *r*_2_ *=* 0.1*r*_0_ so as to prevent branch overlaps and to enable the new segment to grow to *r*_0_ over time. Assumption 6 is integrated into the model by turning off forking after a certain threshold timestep *t*_*B*_ corresponding to the time of the seal’s birth, after which only growth by elongation of branch tips is possible. We choose this birth threshold timestep 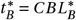, with 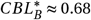 corresponding to the rescaled CBL of our juvenile grey seal specimen (SE1, see below), known to be a neonate (Mason et al., 2020).

### Comparison with experimental data

We compared the structures resulting from this model with the geometry observed in tomograms obtained from grey seal (*Halichoerus grypus*) and harp seal (*Pagophilus groenlandicus*) skull specimens.

Grey seal specimens examined were a neonate (Cambridge Veterinary Anatomy Museum specimen SE1) and an adult (Oslo Natural History Museum specimen 7367), both scanned in the context of an earlier study (Mason et al., 2020). Harp seal specimens examined were three juveniles of various sizes (specimens 7357, 7498, and 7360) and one adult (specimen 7495), all from the Oslo Natural History Museum and scanned in the context of this study, using a Nikon Metrology XT H 225 ST CT scanner with cubic voxel side length 127-300 μm. The processes used for scanning, denoising, and image analysis follow from Mason et al. (2020).

All measurements below are shown as *x±*2*σ* in text, where *σ* is the standard deviation across samples. Quantitative results for the model are obtained as an average over 100 runs of the simulation.

## 3 Results

The structures found in seal maxilloturbinates (Fig. 1) are qualitatively similar to those displayed by the model results (Fig. 3), in terms of spatial distribution, branching patterns and arrangements, intricacy, regions of lower visual density, and meandering path progression. In particular, turning off forking after the gestation time scale 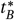 leads to elongated branches at the edges of the system, similar to structures observed in adult seals (Fig. 1d). We did not attempt to show any increases in bone thickness with age, which should be borne in mind when visually comparing model results with tomograms. For quantitative measurements, branch thickness is chosen such that *φ*_model_ ≈ *φ*_obs_ (where *φ*_obs_ ≈ 0.74 obtained from tomograms; this corresponds to the bony structure only, as porosity in a vascularized specimen must be lower, but for simplicity we assume here that the contribution from the mucosa is minimal).

**Figure 3:**
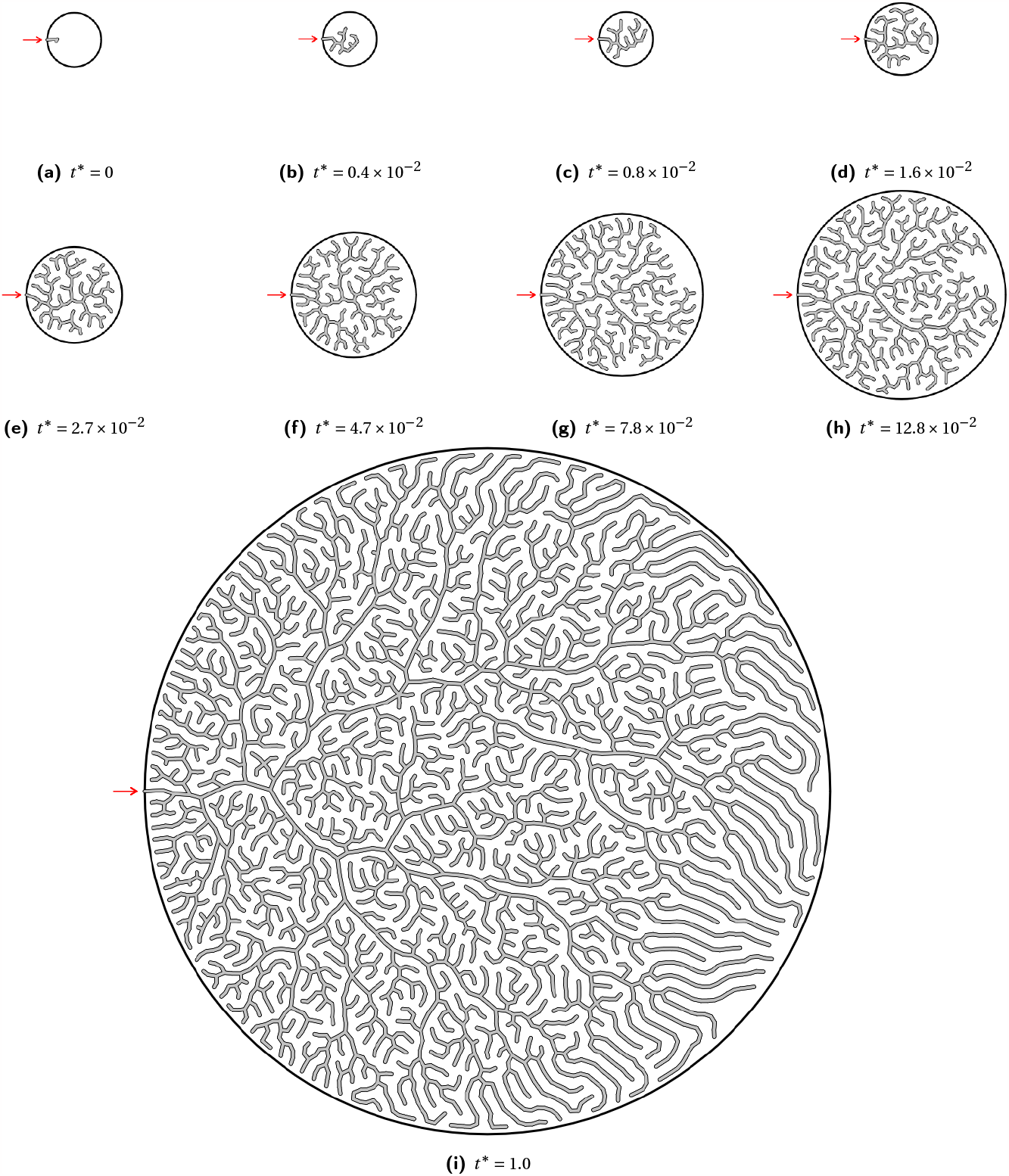
Time series of modelled growth with our model. Boundary circle (nasal cavity walls, with radius *B*) shown as a thick line. The node tree is rooted to the left side of the boundary (red arrows). The final result shows longer branches towards the right side and edges of the system due to forking being turned off and branches only elongating postnatally (i.e. beyond *t* ≥ *t*_*B*_). *t*^∗^ values correspond to rescaled simulation time (i.e. relative age, 0-1).

The condylobasal lengths of the skull specimens are used as a proxy for developmental stage. We define the rescaled condylobasal length *CBL*^∗^ *= CBL*/*CBL*_adult_ ∈ [0, 1] to allow for comparisons between the two species, which otherwise display different condylobasal lengths in adult skulls. The relative age of a seal *t*^∗^, where *t*^∗^ *=* 1 corresponds to a fully developed adult, is defined as *t*^∗^ *= CBL*^∗^ for ease of comparison with simulations. Note that in this definition, *t*^∗^ *=* 0 does not correspond to a seal embryo in its first stage of development, but rather to a theoretical seal whose skull is infinitely reduced.

In order to compare the structures quantitatively, we chose to examine porosity, hydraulic diameter and complexity, which have been measured to compare maxilloturbinates in seals and other mammals in previous work (Mason et al., 2020; Xi et al., 2016), as well as backbone fractal dimension and Strahler statistics, which provide fundamental properties commonly calculated in the geometrical analysis of dendritic and labyrinthine structures (Olsen et al., 2019; Uylings et al., 1975).

### Porosity measurements

Porosities found from the reoriented seal tomograms (without any mucosa attached) are *φ =* 0.71 *±* 0.04 (*n =* 4) in harp seals, and *φ =* 0.74 *±* 0.06 (*n =* 2) in grey seals. Throughout development, the data seem to indicate a slight increase in porosity in grey seals and a slight decrease in harp seals (Fig. 4); these differences likely stem from intraspecific variations in porosity between individual specimens rather than being representative of trends across development in these species. The target porosity in the model was set to *φ =* 0.75 as a direct result of these measurements and therefore remains relatively constant throughout simulation runs (Fig. 4). The actual mean porosity throughout simulations is between 0.74 and 0.76. Root mean squared error (RMSE) between simulated and experimental porosities is *RMSE* ≈ 4.8 *×* 10^−2^ for harp seals, *RMSE* ≈ 7.1 *×* 10^−2^ for grey seals.

**Figure 4:**
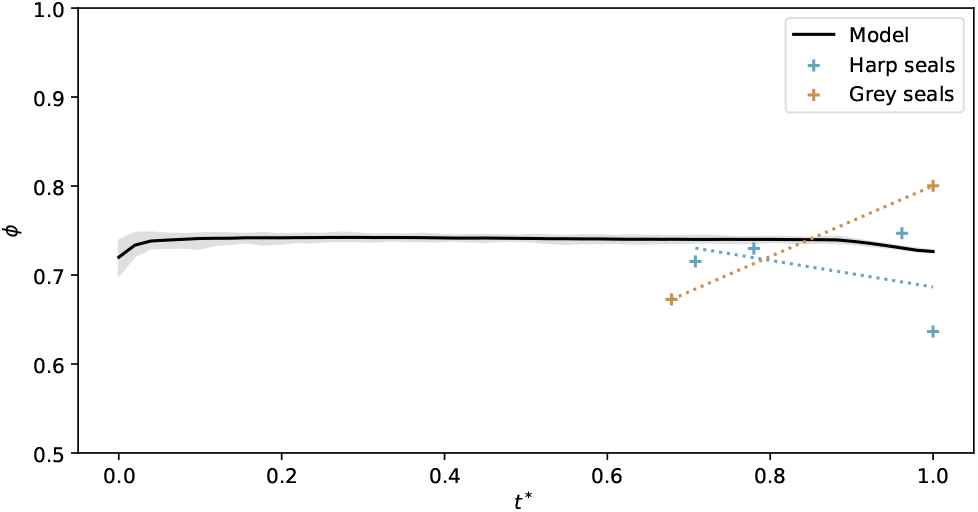
Evolution of porosity (*φ*) as a function of rescaled CBL (relative age; *t*^∗^) measured in seal skulls. Dotted trend lines are fitted via linear regression of actual measurement values in harp seals (blue) and grey seals (red). The black line corresponds to the mean porosity computed through simulation runs, where *t*^∗^ *= t* /*t*_max_ (*t* in relaxation timesteps).

### Hydraulic diameter and complexity

The hydraulic diameter, *D*_*h*_, represents the effective width of air channels, ignoring soft tissue. It is calculated as *D*_*h*_ *=* 4*A*_*c*_ /*P*, where *A*_*c*_ is the total airway area through the chosen reoriented cross-section and *P* the bone perimeter, and has dimensions of length (Mason et al., 2020). We define the rescaled hydraulic diameter as the dimensionless quantity 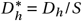, where *S* is the distance between the leftmost and rightmost points on the bone, to be compared with measurements on simulated bone labyrinths. Hydraulic diameters in simulation outcomes display a much lower standard deviation compared with experimental data (Table 1), but the modelled and experimental distributions are consistent. 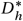 tends to decrease as seals age in both examined species, which the model replicates to a reasonable degree under the *CBL*^∗^ *= t*^∗^ assumption (Fig. 5a) as evidenced by low root mean squared errors: *RMSE*_harp_ ≈ 7.7 *×* 10^−3^, *RMSE*_grey_ ≈ 0.9 *×* 10^−3^.

**Table 1:** Measurements in seal tomograms and model results (mean *±* standard deviation).

|  |  | Harp seals | Grey seals | Model |
| --- | --- | --- | --- | --- |
| Rescaled hydraulic diameter | $D_h^*$ | $(2.2 \pm 0.4) \times 10^{-2}$ | $(2.4 \pm 0.2) \times 10^{-2}$ | $2.5 \times 10^{-2}$ |
| Complexity | $C_x$ | $1.58 \pm 0.02$ | $1.55 \pm 0.02$ | 1.53 |
| Backbone fractal dimension | $d_m$ | $1.18 \pm 0.03$ | | $1.12 \pm 0.04$ |
| Strahler number | $\mathcal{H}$ | $6.6 \pm 0.5$ | | $6.2 \pm 0.4$ |
| Branching ratio | $R_B$ | $3.7 \pm 0.2$ | | $3.6 \pm 0.3$ |
| Branch length ratio | $R_L$ | $2.1 \pm 0.2$ | | $1.8 \pm 0.2$ |

**Figure 5:**
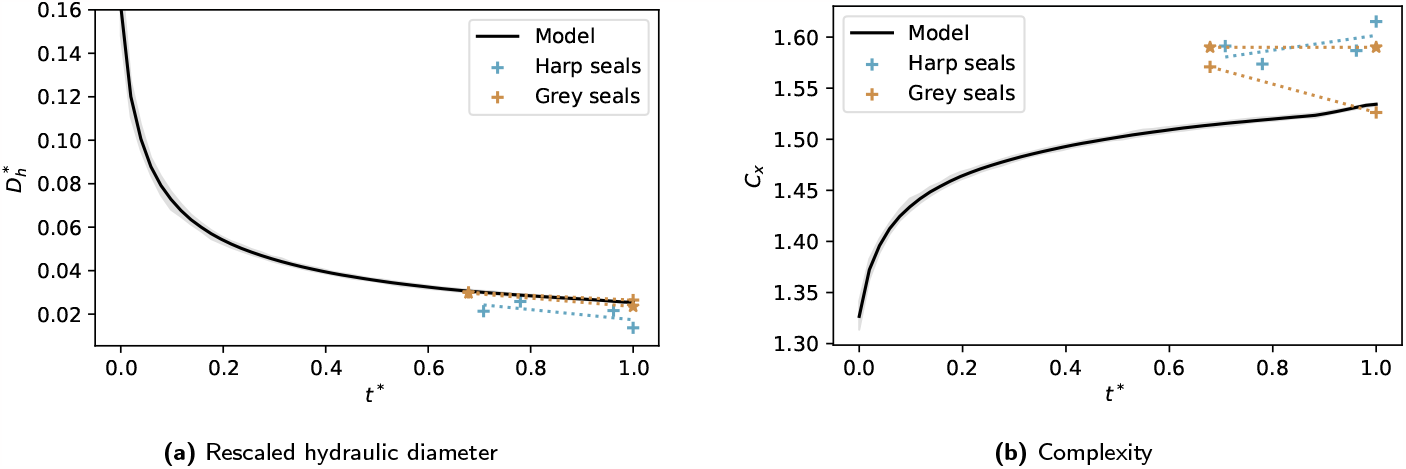
Evolution of the rescaled hydraulic diameter 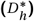 and complexity (*C*_*x*_) in both simulations and seal tomograms as a function of simulation timestep and relative age (*t*^∗^). Grey seal values (orange stars) measured by Mason et al. (2020) (*D*_*h*_ values rescaled by a measured nasal cavity width of respectively 29.0 mm and 45.1 mm).

The complexities (*C*_*x*_ *=* (2 ln *P* − ln(4*π*))/ ln *A*_*c*_, treated as a dimensionless quantity (Mason et al., 2020; Xi et al., 2016)) we measured in grey seal cross-sections are lower than those measured by Mason et al. (2020) (Fig. 5b). Measured complexities are greater in harp seals than in grey seals, consistent with the established relation between complexity and habitat climate across seal species (Mason et al., 2020). Simulation tends to result in slightly lower complexity values (Fig. 5b), which could be attributed to noise levels in scans artificially increasing perceived complexity. *RMSE*_harp_ ≈ 0.07, *RMSE*_grey_ ≈ 0.04.

### Backbone fractal dimension

Plotting bone-bone geodesic distances (i.e. those from one part of the tree to another, walking along the branches) on the backbone tree as a function of Euclidean distances yields the backbone dimension (or path dimension) as the inverse of the slope of the linear fit of the data points (Olsen et al., 2019).

Measuring the backbone dimension requires a representation of the maxilloturbinate cross-section as a connected graph, obtained from the image. While most of the scans are, after denoising, well-connected enough to avoid introducing significant errors (connectivity – ratio of area of the largest connected section of bone to the total turbinate bone area – between 92 % and 99 %), the adult grey seal tomograms are too noisy for the connectivity to reach any greater than 6 %. Therefore, the adult grey seal specimen is excluded from this analysis (and similarly from the Strahler analysis, see below), and we only show results for the harp seal specimens.

The distributions of backbone fractal dimension in harp seals and model results are reasonably similar (Table 1). Relaxing our fourth assumption (that the structure rigidifies over time), by taking *τ* → ∞ in our model, results in early branches stretching as more new branches are added; those branches consequently have a lower backbone dimension.

While the backbone dimension appears to increase slightly as a function of age in seals while not noticeably doing so throughout simulations, the seal tomogram measurements fit reasonably well within bounds of measurements in model results (Fig. 6), with a root mean squared error of *RMSE* ≈ 6.6 *×* 10^−2^.

**Figure 6:**
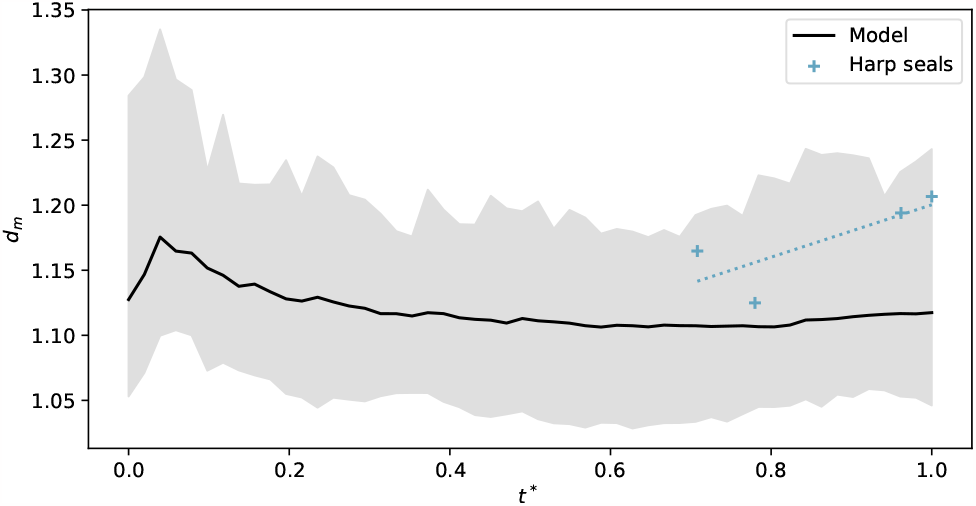
Backbone dimension (*d*_*m*_) as a function of relative age (*t*^∗^). Shaded grey area represents the limits of *d*_*m*_ computed at each timestep over 100 individual simulation runs.

### Strahler analysis

Each of the individual branches in a turbinate cross-section can be assigned a Strahler order following a simple algorithm: leaf branches are assigned order 1 and pruned, leaving new leaf branches which can then be assigned order 2 and pruned, and so on until only the root branch is left (Horton, 1945; Strahler, 1957; Uylings et al., 1975). Colour-coding branches by Strahler order yields Strahler diagrams for the various seal tomograms and model results, which display similarities such as the visual weight of the various colours. High-order (root) branches in seal scans tend to be more winding than in simulation outcomes in which the root branch can appear straighter (Fig. 7). The Strahler number *ℋ* (Strahler, 1957) which denotes the order of the root branch, is consistently between 5 and 7 across both simulations and tomograms (*ℋ*_model_ *=* 6.2 *±* 0.4, *ℋ*_harp_ *=* 6.6 *±* 0.5), with similar distributions and scaling behaviour with respect to rescaled time (*RMSE* ≈ 0.68, Fig. 8a).

**Figure 7:**
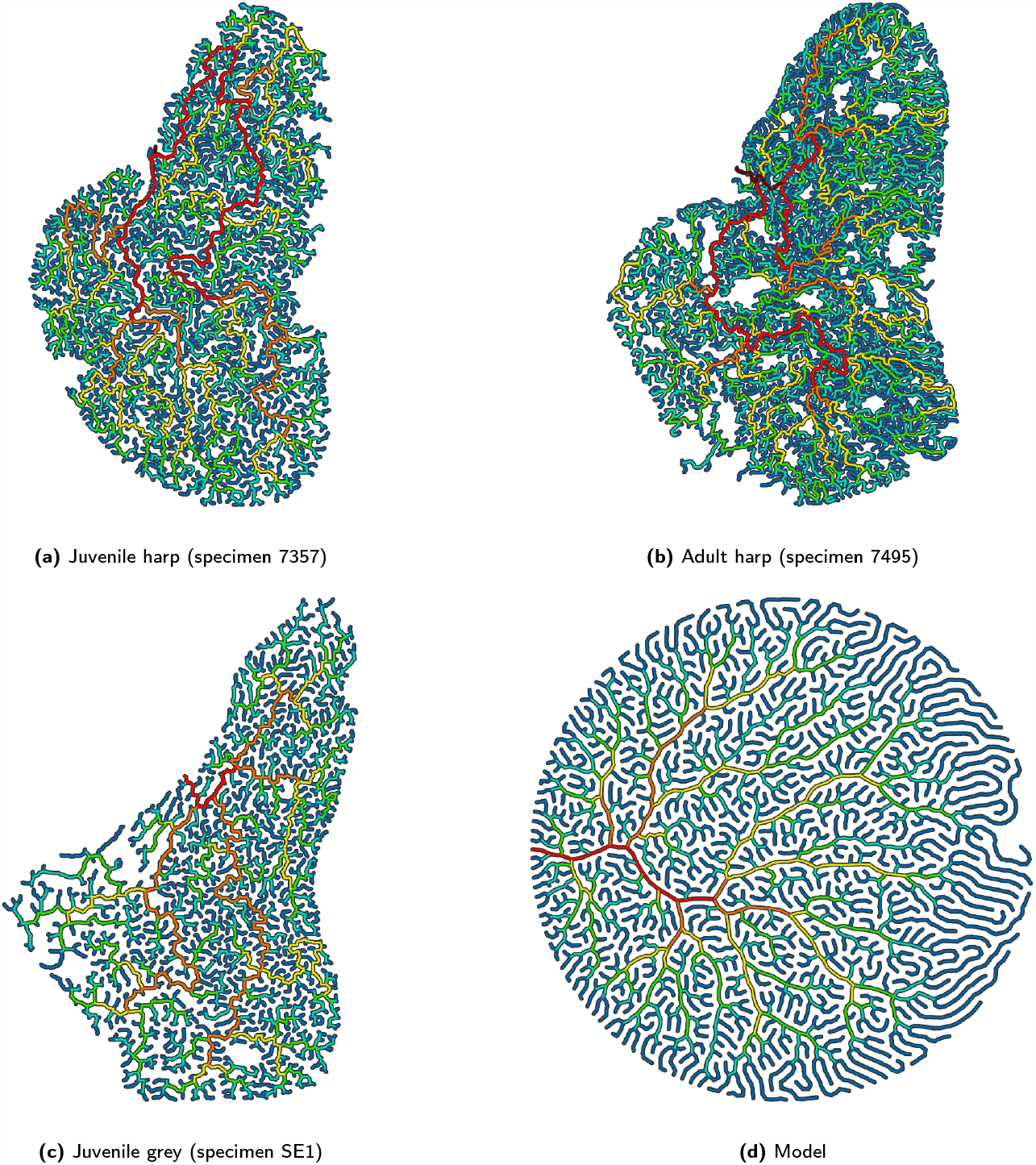
Seal tomogram and simulation Strahler diagrams. The root node (base of the root branch *w = ℋ*, in (dark) red) is selected in tomograms to correspond with the maxilloturbinate root. Branches with low Strahler orders are shown in colder colours (*w*_blue_ *=* 1), with warmer colours corresponding to higher orders.

**Figure 8:**
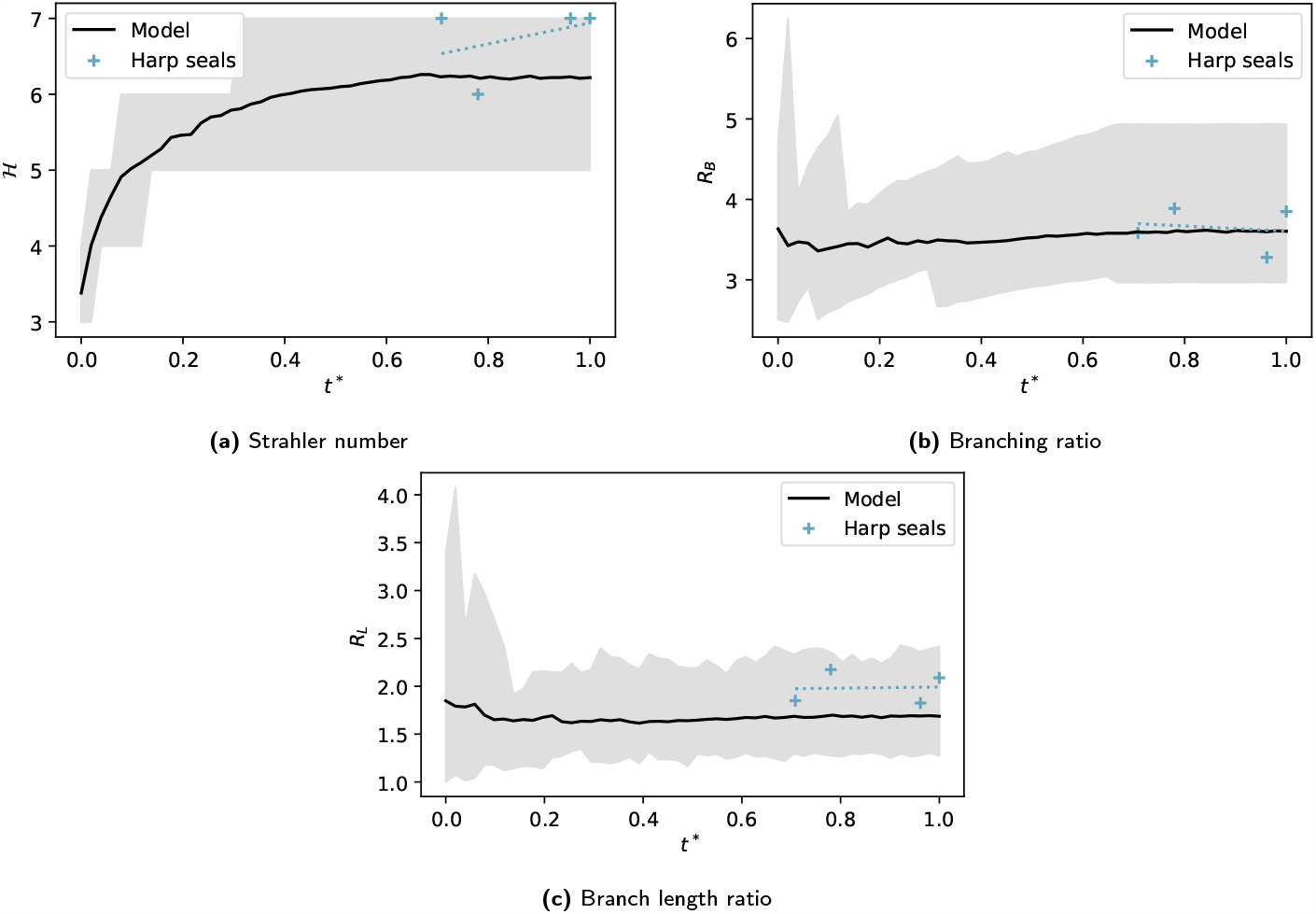
Strahler number (*ℋ*), branching ratio (*R*_*B*_) and length ratio (*R*_*L*_) as functions of relative age (*t*^∗^).

Also examined are the branching ratio *R*_*B*_ and branch length ratio *R*_*L*_, which are given by the slope of, respectively, number of branches and lengths of branches as a function of Strahler order (Barker et al., 1973; Horsfield, 1976; Maxwell, 1955; Olsen et al., 2019). These measurements reflect a geometrical aspect of the system that does not derive from any one parameter in the model – the only way to tweak the model in order significantly to affect the *R*_*B*_ and *R*_*L*_ measurements would be by encouraging growth in high-density areas. While *R*_*B*_ is similar across simulations and tomograms, the length ratio distributions are more disparate (Table 1). Both remain roughly constant as a function of time/age in simulations and tomograms (Figs. 8b and 8c), with root mean squared errors of *RMSE* ≈ 0.25 and *RMSE* ≈ 0.33, respectively.

### Model generalizability

Measurements made in specimens at different developmental stages and through simulation runs show that model timesteps and relative specimen ages (using the index of condylobasal length) are approximately linearly related (Figs. 5a, 6, 8) with relatively low root mean squared errors. To test the generalizability of the model, we ran simulations with the greater porosity *φ*_monk_ ≈ 0.80 observed in Mediterranean monk seal (*Monachus monachus*) maxilloturbinates as the target porosity (Fig. 9). These yield results that closely match measurements made on monk seal tomograms (scans from Mason et al., 2020): a greater hydraulic diameter and lower complexity than in harp or grey seals, but similar Strahler statistics and backbone dimension.

**Figure 9:**
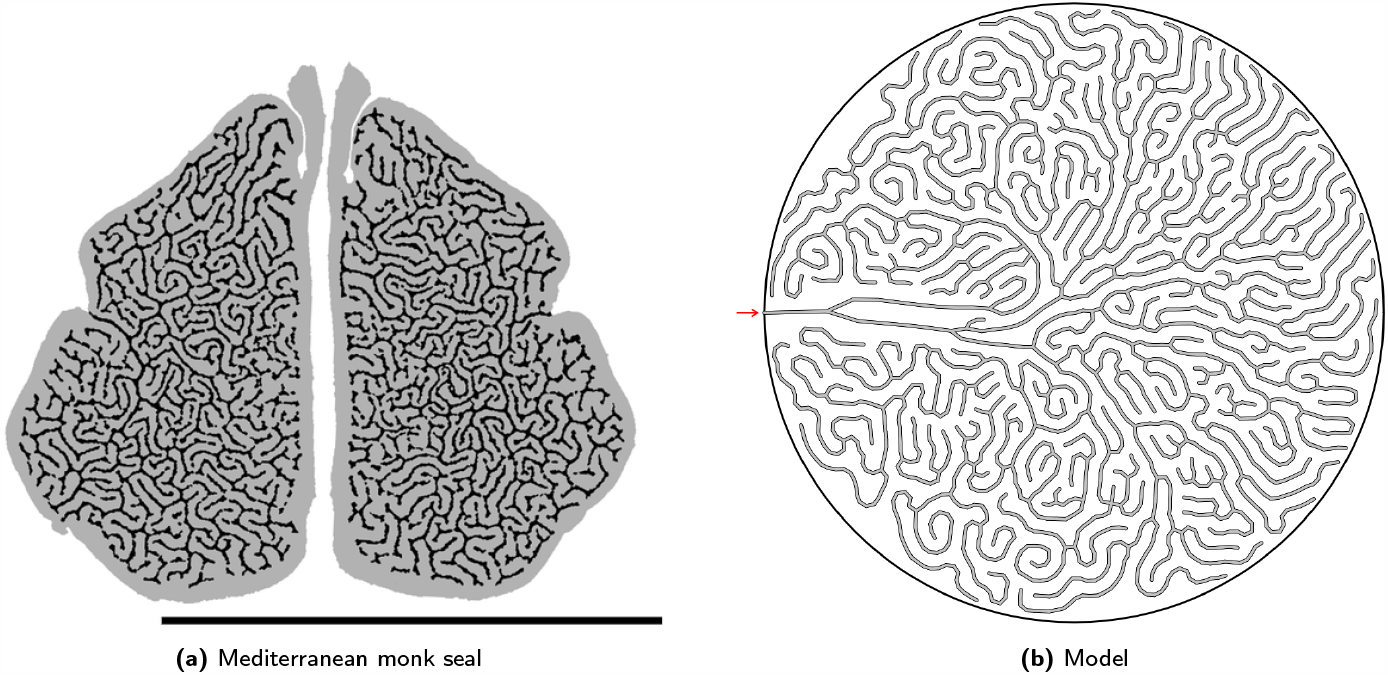
Modelled development of a Mediterranean monk seal maxilloturbinate. Monk seal (specimen NHMO M115) tomogram reproduced from Mason et al. (2020) under the terms of the Creative Commons CC BY license (scale bar 50 mm), juxtaposed to final model result obtained for a target porosity *φ*_monk_ *=* 0.80.

## 4 Discussion

Overall, our results suggest that the rules (assumptions 1-6) embedded within our model yield results comparable with actual seal anatomy, and so may well correspond to biological mechanisms controlling turbinate morphogenesis within seals. This model thus allows us to formulate specific hypotheses concerning how the intricate, labyrinthine pattern may develop.

Assumption 1 of turbinate growth within a confined space is consistent with anatomical observations in other mammals, where the nasal cavity and the turbinate expand in size hand-in-hand, excepting early stages where the main branches grow within a relatively much larger chamber (e.g. Rowe et al., 2005). In our model, this spatial confinement does not affect the key features of the pattern and the avoidance of branch growth into the boundaries can come from a similar biological mechanism as that between branches (assumption 5, discussed below).

Assumption 2, relating to tip growth and forking, was designed to be consistent with observations in other mammals (Martineau-Doizé et al., 1992; Smith et al., 2021). Although bone resorption has been observed in the development of the scroll-like turbinate system of pigs, contributing to the curvature (Martineau-Doizé et al., 1992), seals have a dendritic turbinate pattern and resorption was not considered within our model. The progressive ossification of the maxilloturbinates from root to branch (Smith et al., 2021) will stiffen the branches over time, fixing the initial curvatures (assumptions 3 and 4). In reality, branches nearer the root of the maxilloturbinate tree tend to be thicker than terminal branches, this being more obvious rostrally and caudally in the tomograms, supporting the notion of progressive stiffening. The separation of the branching stage and no-branching stage as prenatal and postnatal (assumption 6) was initially suggested by Mason et al. (2020) based on comparison of grey seal tomograms and is supported here as juvenile harp seal samples show a similar branching depth and number as adults (Figs. 6 and 8b). One notable model implementation that controls the selection of the branching site is that it depends on density – a new branch will more likely appear in a region of low bone density than in a region already packed with branches. Without this constraint, the model yields uncharacteristic, uneven growth (Fig. 10a). The ability of branching points to detect the density in neighbouring areas could be derived from a similar mechanism to the avoidance mechanism (assumption 5).

**Figure 10:**
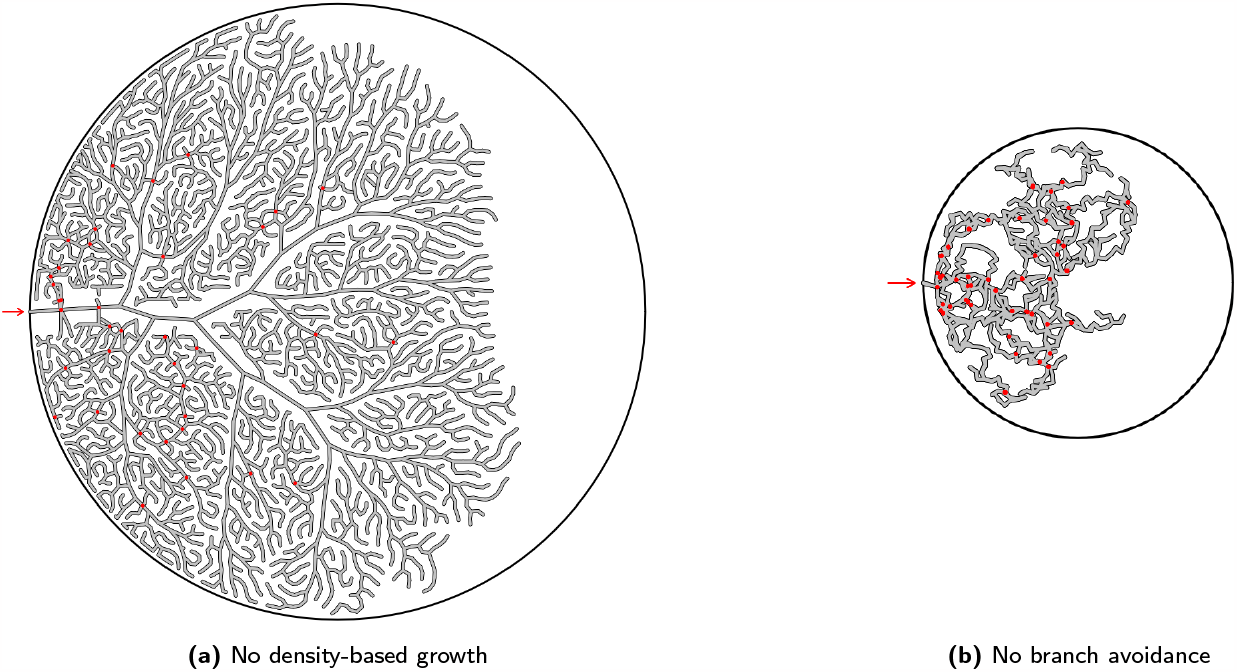
Unstable model variants. Simulation outcomes obtained for two variants of our model; the removal of density-based growth and the removal of branch avoidance. Red dots indicate intersections between non-neighbour branches. Simulations were stopped beyond 50 branch intersections for clarity.

A key assumption of the model for which we have no direct evidence relates to branch avoidance (assumption 5). In the absence of an avoidance mechanism, appositionally growing lamellae would inevitably collide with each other (Fig. 10b). Such a mechanism is consistent with the turbinate growth patterns observed in short-snouted dog (*Canis familiaris*) breeds, where insufficient space in the nasal cavity leads to turbinates veering into the pharynx (Oechtering et al., 2016). The biological basis of this potential avoidance mechanism is currently unknown. How could a branch detect the presence of others across a length scale of hundreds of microns? Following the principles of Occam’s razor, we would prefer to identify one feedback process suitable for guiding maxilloturbinate growth both prenatally and postnatally. There are unlikely to be any significant temperature gradients within the maxilloturbinate mass prenatally, given that the fetus is within the amniotic fluid at maternal core body temperature. A diffusing chemical signal seems unlikely postnatally, given the flow of gas through the turbinate system once the seal begins breathing air, which would sweep away any secreted morphogen. We therefore turn to considering mechanical signals for feedback purposes, which might operate both before and after birth. Turbinates are likely to be subject to shear stress in the longitudinal direction from amniotic fluid prenatally (Koos & Rajaee, 2014) or air postnatally. Blocking of postnatal airflow on one side can affect turbinate development in mice (*Mus musculus*) (Coppola et al., 2014). Coppola et al. estimated shear stress values in the mouse nasal cavity to exceed 1 Pa in some rostral positions, declining significantly towards the olfactory region posteriorly. They proposed that shear stress could be used as a mechanical signal to influence turbinate growth. We hypothesize that cells within the growing branches sense these shear stresses and respond by adjusting proliferation or growth rate. In a channel section of dimensions *ℓ× h × b* (Fig. 11), the shear stress *σ* on the walls in the direction opposite the pressure difference ∆*P* which is driving flow can be computed from the force balance 2*σS = A*∆*P*, with *A = bℓ* the cross-sectional channel area and *S = hℓ* the wall area. This yields 2*σ =* (*b*/*h*)∆*P* ; since ∆*P* is the same in neighbouring channels, it follows that *σ* ∝ *b*, i.e. wider channels have a larger shear stress on the walls. In such larger channels within the maxilloturbinate mass (either between neighbouring branches or between a branch and the confining boundary), the greater shear stress may be sensed either by the bone directly (such as has been described by Tomlinson et al. (2020)) following deformation of the overlying mucosa, or indirectly via surface structures such as cilia (whose role in mechanosensing has been discussed by Wachten and Mill (2023)). Cilia are found on the pseudostratified columnar cells of the seal maxilloturbinate respiratory epithelium (Folkow et al., 1988). In the latter case, downstream cellular signalling involving different cell types would ultimately encourage local bone growth, resulting in space-filling by tip growth and new branch formation. Conversely, in narrow channels (i.e. higher-density areas), the shear stress becomes smaller with the flow and stops stimulating (or even starts inhibiting) cell growth, leading to the stabilization of the narrow channels and the prevention and branch collision and fusion.

**Figure 11:**
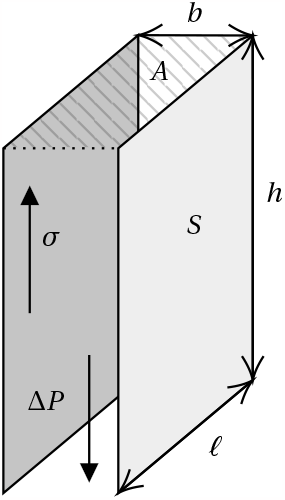
Simplified geometry of a channel of dimensions *ℓ×h×b* with pressure gradient ∆*P* and shear stress on the walls *σ*.

Given such an influence of shear stress on bone growth, the overall porosity of the turbinates can become an optimized target for individual species to achieve through development, and a trait evolutionally tunable by the genetic factors that regulate the gain in the mechanosensory system. This will produce the observed relatively stable porosity in a seal’s nose throughout its postnatal development, which is variable between species but important for the heat and water exchange of the animal (e.g. Cheon et al., 2023; Folkow and Blix, 1987). Testing our hypothesis will require experimentation, for example altering the mechanical stresses experienced by the turbinates during prenatal growth, and then examining the genetic and cellular responses of the tissue. These experimental tests may be possible in a more accessible carnivore model, such as canine or feline, in which the turbinates likely share similar patterning mechanisms albeit having a less complex pattern (Van Valkenburgh et al., 2004). Direct tests on the seals would best be approached with an in vitro culture model, where turbinate branch tip cells are grown and differentiated within controllable fluid environments.

## Data availability

Raw tomograms and images from simulations are available from the authors upon request.

## Code availability

C++ model implementation code and Python analysis code are available from the authors upon request.

## Acknowledgements

We thank Arnoldus Schytte Blix and Léa Wenger for helping to produce some of the CT scan data used in this study. This work was partly supported by the Research Council of Norway through its Centers of Excellence funding scheme, project number 262644.

## Author contributions

Ø.H. produced the harp seal CT scans. All authors took part in discussions and defining the biological basis for the model; J.E.K. and E.G.F. implemented the technical aspects of the model. J.E.K. performed the numerical simulations and analysed the data. M.J.M., F.X. and L.P.F. developed the biological theory and discussion. All authors contributed to writing and proof-reading the paper.

## Materials & correspondence

Correspondence and requests for material should be directed to Jonathan E. Kings.

